# Pathways Analyzer: Design of a Tool for the Synthetic Assembly of Escherichia Coli K-12 MG1655 Bacteria for Biofuel Production

**DOI:** 10.1101/634998

**Authors:** Laura Vasquez, Ricardo Alvarado, Allan Orozco

**Affiliations:** Department of Computer Engineering, Costa Rica Institute of Technology (TEC), Cantoblanco, Madrid, Spain; National Laboratory of Nanotechnology (LANOTEC), Nanobiotechnology Department, Cantoblanco, Madrid, Spain; School of Health Technology, Faculty of Medicine, University of Costa Rica (UCR), Cantoblanco, Madrid, Spain; Department of Molecular Biology, Faculty of Science, Autonomous University of Madrid, Cantoblanco, Madrid, Spain

## Abstract

**Summary:** Due to the impact of environmental pollution, the importance of producing high quality biofuels and to leverage organic waste that normally would have no use has increased over time. Through synthetic biology, it is possible to improve existing organisms to process waste that is traditionally not used for biofuel production, such as whey.

With the redesign of metabolic pathways, it is possible to create connections for the implementation of new organisms that carry out functions that are normally not present in nature.

From a computational point of view, metabolic pathways, which can be found in data sources as KEGG, can be converted to a graph data structure. These transformations enable the use of well-known algorithms, which enables the optimization of the analyses required to achieve the assembly of new organisms.

The present work aims to design a tool for the transformation of metabolic pathways and the development of path finding algorithms that establish relevant links between compounds that are essential to the biofuel production process.

As a result, a catalog of biobricks is created from the analysis of a subset of paths which can be used in the design stage of the synthetic assembly of the *E. coli* bacteria. The assembly’s structure and functions are characterized according to the pieces used.

Finally, new constructions are visualized with the goal of demonstrating and supporting the analysis processes, thus assisting people that work in the field of Synthetic Biology.

**Availability:** Pathways Analyzer is accessible at: https://gitlab.com/lvasquezcr/pathways-analyzer/

## 1 INTRODUCTION

Synthetic biology is a new science that enables the design and construction of new genetic components, devices and systems that will carry out new functions that are usually not found in nature, through the reprogramming of their genetic code (Synthetic Biology Community, 2013).

When a new genetic element is created, a standardization process is performed abstracting its composition through a connection interface that reduces its complexity. This helps organizations that work in organisms’ synthetic assembly to share knowledge between them. Through this concept, the assembly of a new organism is simplified through blocks or “biobricks” making this process equivalent, at a high level, to put together a puzzle (Heinemann & Panke, 2006).

Through synthetic biology, it is possible to modify organisms so that they participate in the degradation process of organic matter for biofuel production, providing better alternatives for biofuel sources. Recent studies show that the reengineering of the *Esche-richia coli* (*E. coli*) bacteria allows for the production of many types of biofuels (Chin, 2008) and that it is possible the production of renewable biofuels, through the modification of the metabolism of this bacteria (Howard, Middelhaufe, & Moore, 2013).

In nature, many organisms work as small factories, meaning that from raw material, subproducts are generated and used for other particular purposes. Each process within these organisms is called a metabolic pathway, which illustrates how primary components are degraded into more simple compounds until they reach the goal product (IvyRose Ltd., 2014). For example, the *E. coli* bacteria can be modified to process organic waste such as whey into fatty acids for biofuel production (Liu & Khosla, 2010).

The Figure 1 shows how whey, also called milk serum, is used as a primary compound to produce biofuels. Proteins break down source compounds into simpler compounds that will be essential in the process of biofuel production. Later in the process, the genes that produce the relevant proteins will be identified and then transformed into the standardized blocks. The design stage of metabolic pathways is complex since when there are many possible options of how these can be modified.

**Fig. 1.**
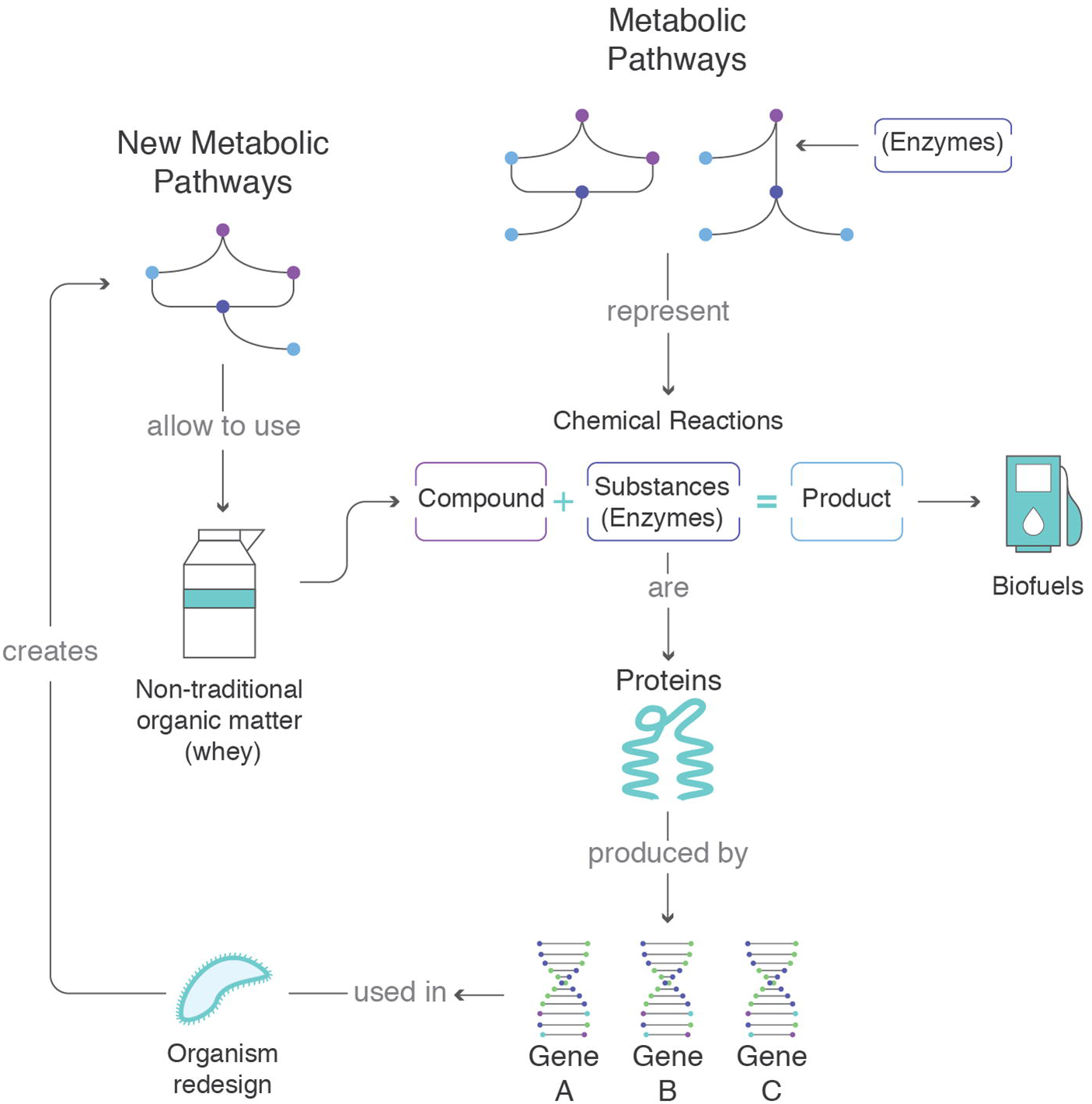
Design of new metabolic pathways

Current tools as SynBioSS (Hill, Tomshine, Weeding, Sotiropoulos, & Kaznessis, 2008), GeneDesigner (Villalobos, Ness, Gustafsson, Minshull, & Govindarajan, 2006) and GeneComposer (Lorimer, Raymond, & Walchli, 2009) provide a generic assembly of biobricks with no guidance. Moreover, there are not many Bioinformatics tools available for biofuel production that assists researchers with the assembly of new genetic components, suggesting possible genetic pieces and describing the final product structurally and functionally.

The present project Pathways Analyzer, is a tool that aims to provide guidance to researchers with the optimization of metabolic pathways and the construction and filtering of biobricks. These biological pieces will be further used for the synthetic assembly of the *E. coli* genome in the process of organic matter (whey) degradation. Also, it will improve the efficiency of the traditional process to design metabolic pathways, in the transformation of components and products, through the incorporation of bioinformatics elements in this design stage. This tool will contribute to the automation of the analyses on the gene selection for the design of new metabolic pathways.

## 2 PROTOTYPE SOURCE DATA

Entries to carry out the prototype’s tasks were delimited according to the project’s scope. For metabolic pathways analysis, a group of start and end components was selected based on the purpose of biofuel production. The initial nodes were selected depending on their existence in organic waste ideal for this purpose. Lactose was chosen as the main input of the analysis since it is present in whey. Terminal nodes were part of the energy producing process in many organisms, which have an important role on biofuel production. Based on this, Pyruvate was chosen as the main end node.

Metabolic pathways repository was chosen based on the protocol for data access, output formats, libraries available for the transformation of metabolic pathways, efficient access to the gene and protein references in other databases and local expertise. This way KEGG (Wrzodek, Dräger, & Zell, 2011) was selected as the main data source for metabolic pathways for their immediate translation of metabolic pathways to a standard format on the computing environment. Maps selections were limited to the organism *Escherichia coli K-12 MG1655*.

KEGG entries has direct references to external data bases such as the Universal Protein Resource (UniProt Consortium, 2014), which has a simple programming interface and great relevance for scientific community in the genomic field.

For the biobricks assembly, the plasmids (pB1A3, pSB1T3, pSB3K3) and restriction sites (EcoRI, PstI, XbaI, SpeI) were used, based on what iGem Foundation (International Genetically Engineered Machine Foundation) defined as a standard.

The information about the proteins and genes involved will be gathered once the required metabolic pathways were selected, this will enable a catalog for the synthetic assembly of the *E. coli* bacteria. The catalog will be built under the following sources:

1. The iGem Catalog from the Standardized Biological Parts Registry (International Genetically Engineered Machine Foundation).
2. A preselected list of products obtained from the analyses of the tool (International Genetically Engineering Machine Foundation).
3. Custom sequences added by the user.

## 3 PROTOTYPE DEFINITION

### Tool Design

Before the development of the prototype, a computational model was defined to organize all the components involved in this process. The design is divided in seven stages which encompass the inputs collection and the metabolic pathways analysis to the assemblies’ construction and visualization.

The Figure 2 shows in detail the different stages that will be described below:

**Fig. 2.**
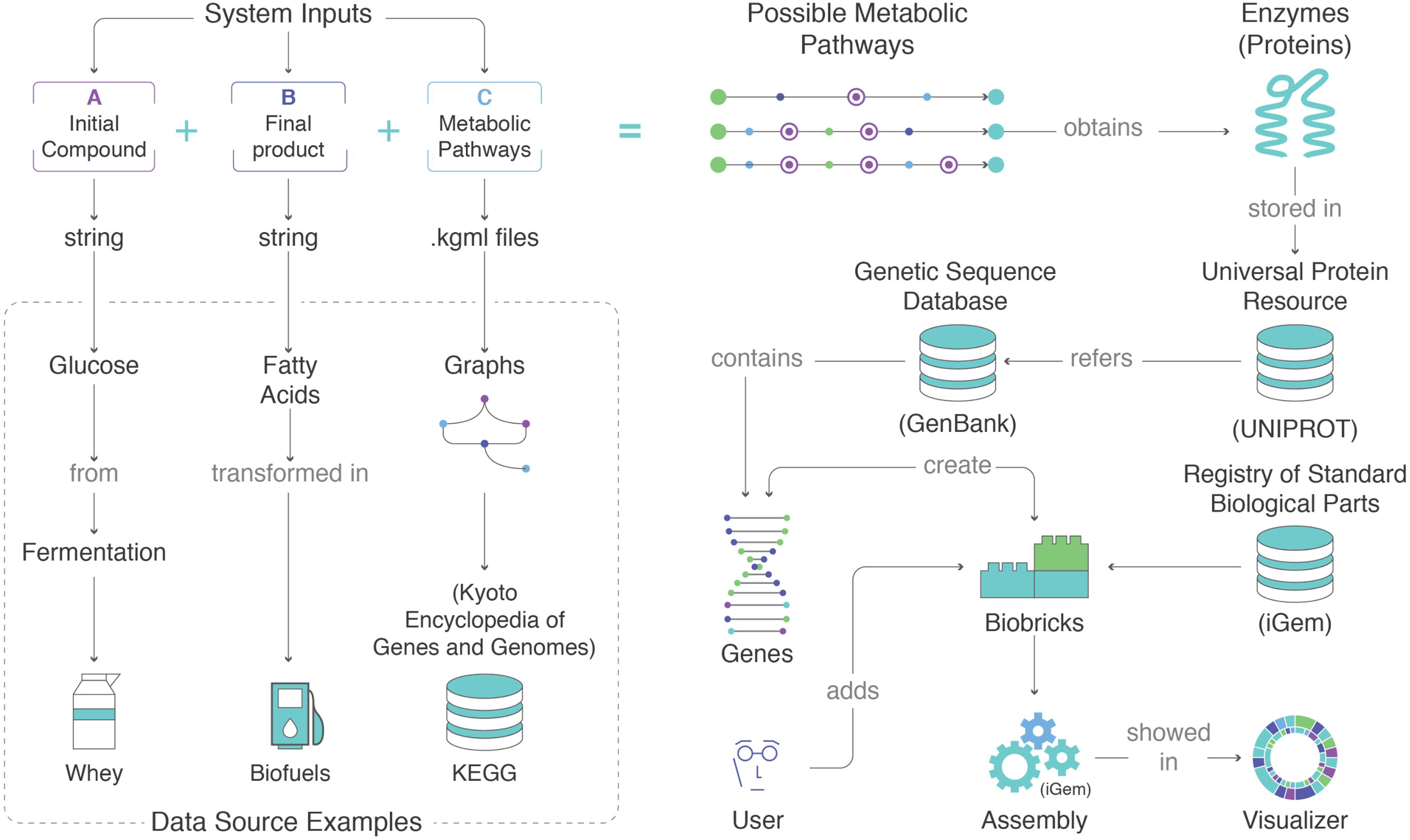
General procedure of data analysis

1. *Entry identification:* the initial and final nodes are selected with the purpose of finding the possible paths between both. These searches will be limited by depth and restrictions defined by the user as the inclusion and exclusion of components. Start compounds will belong to the carbohydrates group (sugars), end compounds to the fatty acids group, such as triglycerides. Maps from KEGG database will be extracted in its standard KGML format.
2. *Metabolic pathways calculation:* the initial and final nodes are submitted, searches on KEGG will be performed to calculate the possible paths existing over the available maps.
3. *Paths selection:* each route will be evaluated by its length and the number of enzymes involved. The user will select the best route according to the needs of the corresponding study.
4. *Proteins and genes selection:* once the best route is selected, the enzymes that are involved in the process will be extracted and the related information will be retrieved from UniProt. This database contains references to GenBank, where the user will get more data about the genes responsible for this proteins’ production.
5. *Biobricks Creation:* using the genes from the previous step, new biobricks will be created, that later will form the proposed catalog.
6. *Creation of an assembly:* after providing the required biobricks, they will be loaded in a catalog and the user will create assemblies for *E. coli* bacteria, according to the rules defined by iGem.
7. *Assembly characterization and visualization:* once the assembly is built, it will be characterized in a structural and functional way, with regards to the proteins involved in the process.

### Prototype implementation

Pathways Analyzer is a command line application written primarily in Java. Support tasks were developed in Ruby for the elimination of nested pathways and Perl to obtain complementary information from GenBank database. Visualization of pathways was achieved through JUNG, the Java Universal Network/Graph Framework (The JUNG Framework Development Team) and assemblies with a simple Javascript webpage.

The redesign of metabolic pathways required a deep analysis in databases such as KEGG. There were different ways of breaking down one or more components into another and some of these ways were better depending on the goal or the purpose of the analyses. Some paths were longer or shorter, depending on the number of components and proteins involved. Proteins will be the most relevant since they carry out chemical reactions to catalyze these decompositions and are part of each organism’s metabolism. However, before selecting the required proteins, and thus the genes involved for the organism’s reengineering, the pathway that adapts best to the analysis requirements was chosen.

In the first stage of the metabolic pathways analysis and selection, all the inputs were collected. The KEGG Pathways database was queried through its webpage to extract all the metabolic pathways that matched with the keywords "Lactose" and "Pyruvate" (Kanehisa Laboratories, 2018). Once the list of required maps was compiled, they were downloaded from UNIX using Ruby.

Next stage was the translation of KEGG maps from KGML to GraphML format. The latter enabled the parsing of metabolic pathways into a format that could be easily translated into a standard data structure for computing data. KEGGtranslator (Wrzodek, Dräger, & Zell, 2011) was used to make this translation, and from all of its available output formats, GraphML was chosen due to its compatibility with the subsequent libraries used on the implementation of this prototype. However, not all the maps went through a straight-forward translation, since some of them had an unsupported structure for the KEGGtranslator.

Due to the nature of KGML files, it was common to find clusters of enzymes in the breakdown of one component to another shown. This condition made the translation library crash and since there was no support regarding this issue, it was necessary to go through a workaround. As a solution, a preprocessing script in Ruby was developed, loading these files into a flexible XML object that allowed fixing the inner structure of the graph.

The result was a consolidated graph that will be used for pathfinding algorithms in order to search for the most relevant results for the next stage. Merging all the maps this way enabled new connections between the different nodes that didn’t exist before, which left the opportunity to find new results on the design of metabolic pathways. The consolidation of all the inputs into an integrated network enables new paths, such as the path from A → E in Figure 3.

**Fig. 3.**
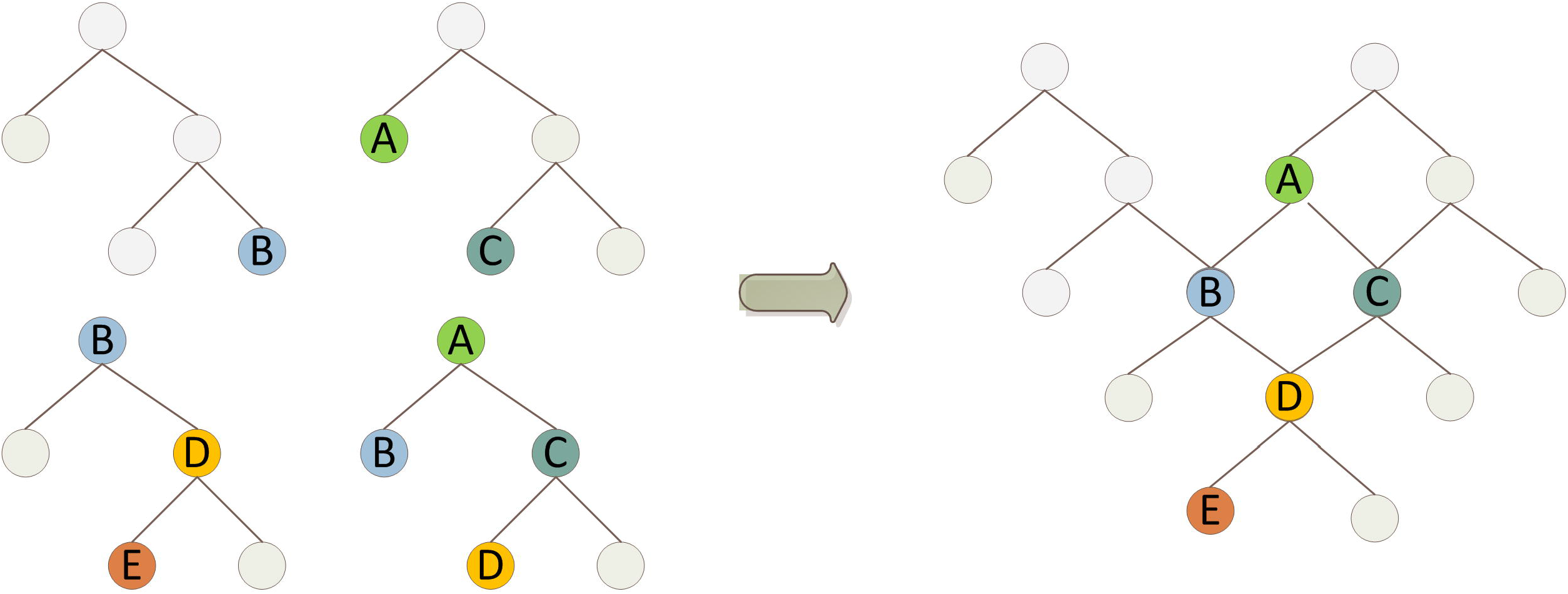
Union of independent networks into one only structure

In order to find new paths between two nodes, pathfinding algorithms such as Dijkstra (Morris, 1998) and Breadth First Search BFS (Weisstein, 2014) were included in the prototype. The shortest path between two nodes was achieved using Dijkstra algorithm which was built into the JUNG library. The library’s available capabilities for modeling and visualization of data also enabled later the graphical representation of results.

It was also necessary to find all the available paths between two different nodes. The resulting analysis determined which of them was more relevant; this was not necessarily given by the shortest distance. To find all the available paths between two nodes the algorithm was implemented based on a restriction system. This system limited the algorithm scope in terms of depth, required and excluded compounds, according to the user’s requirements. This will filter and prioritize the results depending on the inputs given. In terms of results visualization, it was relevant to know which proteins were involved in the final path and the side compounds derived from the breakdown of more complex compounds, catalyzed by the present proteins. Even though these components were not part of the critical route, the user had to be aware of all the subproducts produced in a reaction. Based on this information the results will be visualized using JUNG, through a color label system. Proteins were highlighted in green; the components derived but that are not a part of the path in gray and the critical route in red, as in Figure 4.

**Fig. 4.**
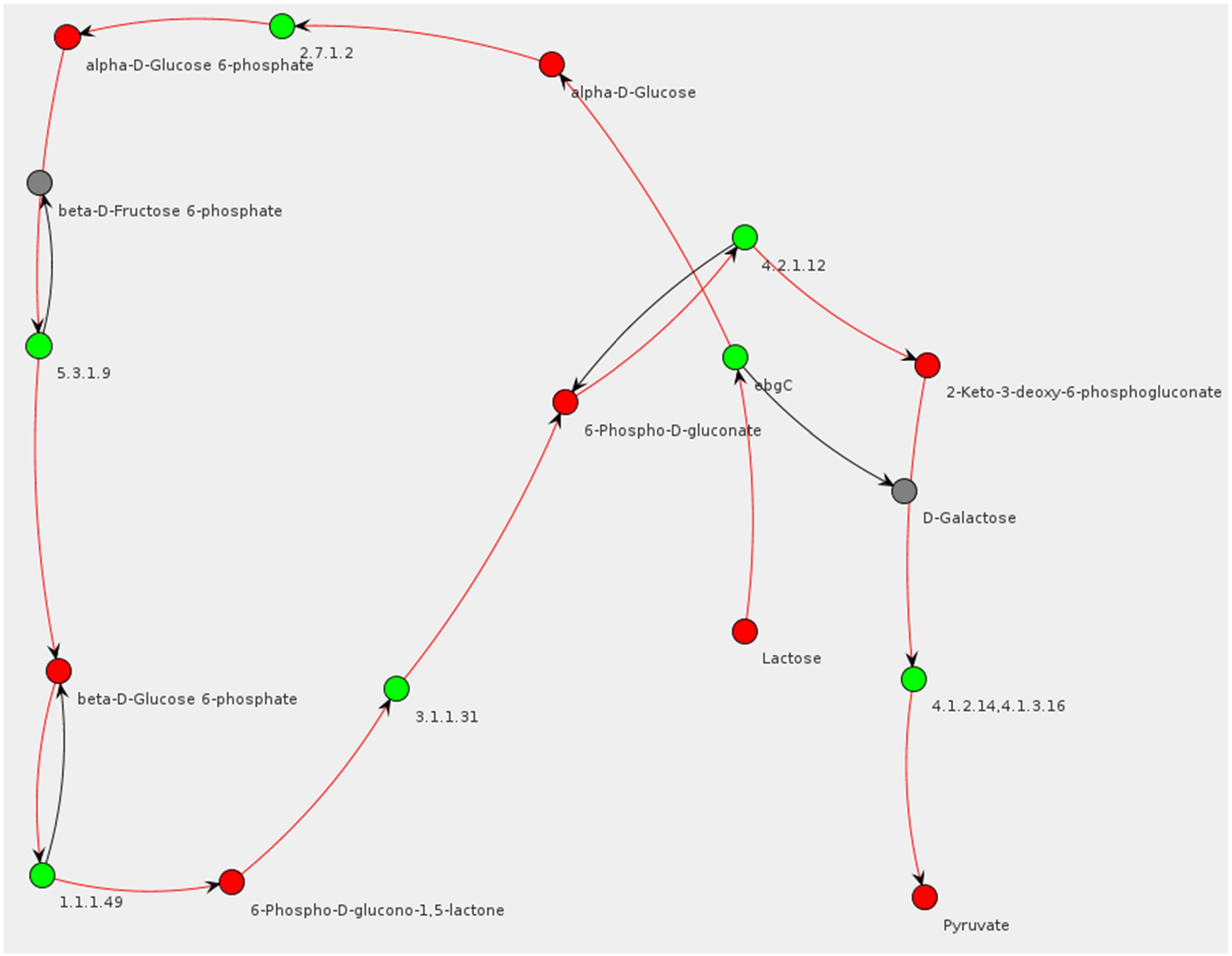
Visualization of the shortest path between Lactose and Pyruvate

In the next stage, algorithms were based on the selected path from the available information gathered from its proteins. The information extracted from the KEGG database had a direct connection from its proteins to UniProt, a data source of protein sequences and functional information. The main advantage was that it had a programming interface that allowed the data extraction through the REST API (UniProt Consortium) in a variety of formats. This was achieved thanks to the UniProt ids that KEGG provided, to obtain further information about them, relative to the genes that produced them, the organisms they belong to, their origin and mainly the cross references to other databases, such as GenBank.

Proteins can be encoded in nature from different types of genes and each of these genes may be a key piece for the assembly of new organisms. To carry out this task, GenBank (Benson, Cavanaugh, & Clark, 2012), a database of genetic sequences, was selected to query additional information of each gene using the “Entrez Programming Utilities” (National Center for Biotechnology Information, 2018).

In particular, it was possible to extract the coding regions of the genes, that is, the sequence of instructions to produce the protein, commonly known as DNA. Using “Ebot”, interactive web tool to personalize and generate scripts based on a sequence of steps, it was possible to retrieve the necessary information from these databases to form the basis of the biobricks.

After proteins and genes were analyzed, the creation and validation of a catalogue with the required components for the synthetic assembly of the *E. coli* bacteria became possible.

The inputs related to metabolic pathways were extracted from *Escherichia coli K-12 MG1655*. The catalog of biobricks (Figure 5) will come from three possible sources: genes as a result of the explained analysis, existing biobricks in the Registry biobricks iGem or customized biobricks built by the user.

**Fig. 5.**
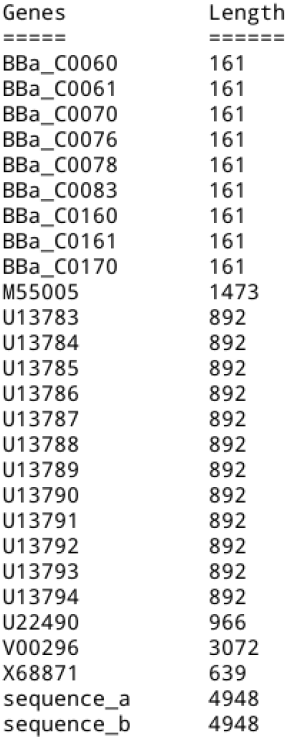
Biobricks Catalog

This catalog of biobricks allowed for the design of new organisms for biofuel production, through the improvement of their metabolic pathways. The obtained assemblies were defined by the standard already proposed by iGem. Once the catalog was defined, it was possible to follow a methodology where the biobricks are assembled onto plasmids or between them, with the purpose of verifying its compatibility, between them (International Genetically Engineered Machine Foundation, 2019).

As the final step, the assemblies built were characterized as follows:

1. Structural characterization: measured the length and the amino acid number.
2. Functional characterization: stated an approximation of the functions that the final assemblies may have. This feature is available when the inputs are obtained from the analysis of metabolic pathways. In this case, the inputs’ source is known and the relevant information about its possible functions is available.

The developed prototype was run with 101 metabolic pathways corresponding to all the maps that matched Lactose and Pyruvate as compounds. Analyses were carried out to a depth of 12 nodes, over a graph of 4,173 nodes and 5,773 edges. After the analysis of 229,753 possible connections 14 pathways were found.

Moreover, with the purpose of refining these searches, some compounds were excluded. When the compound “alpha-D-Glucose” was added to the exclusion list, analysis towards 14 nodes of depth was performed. As a result, 10 connections between the original compounds were found after carrying out an analysis of 78,323 pathways. On the other hand, the inclusion of the compound “D-Galactose”, resulted in a 15-depth nodes search finding 785 matches over 1,670,100 analyzed pathways. At the end, after applying this filter, the list of results was reduced to 47 pathways found.

In Figure 6 is shown how the performance increases when a set of exclusions is added to the algorithm. Although, additional searches were done to a higher depth, the amount of pathways found was similar.

**Fig. 6.**
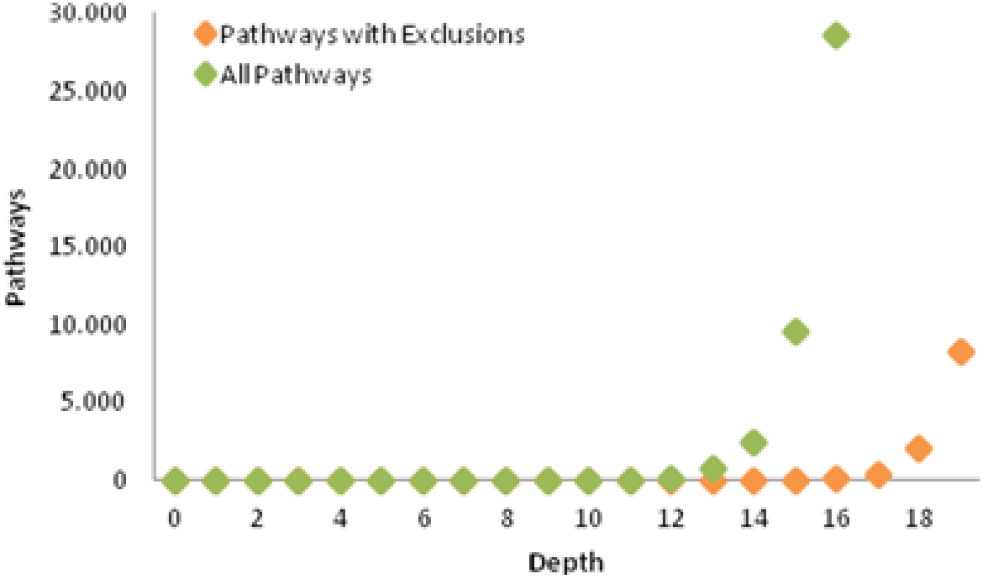
Performance increase with exclusions

Finally, with the goal of showing the interaction between the selected genes and plasmids, the finished assemblies are visualized in a web page, as shown in Figure 7.

**Fig. 7.**
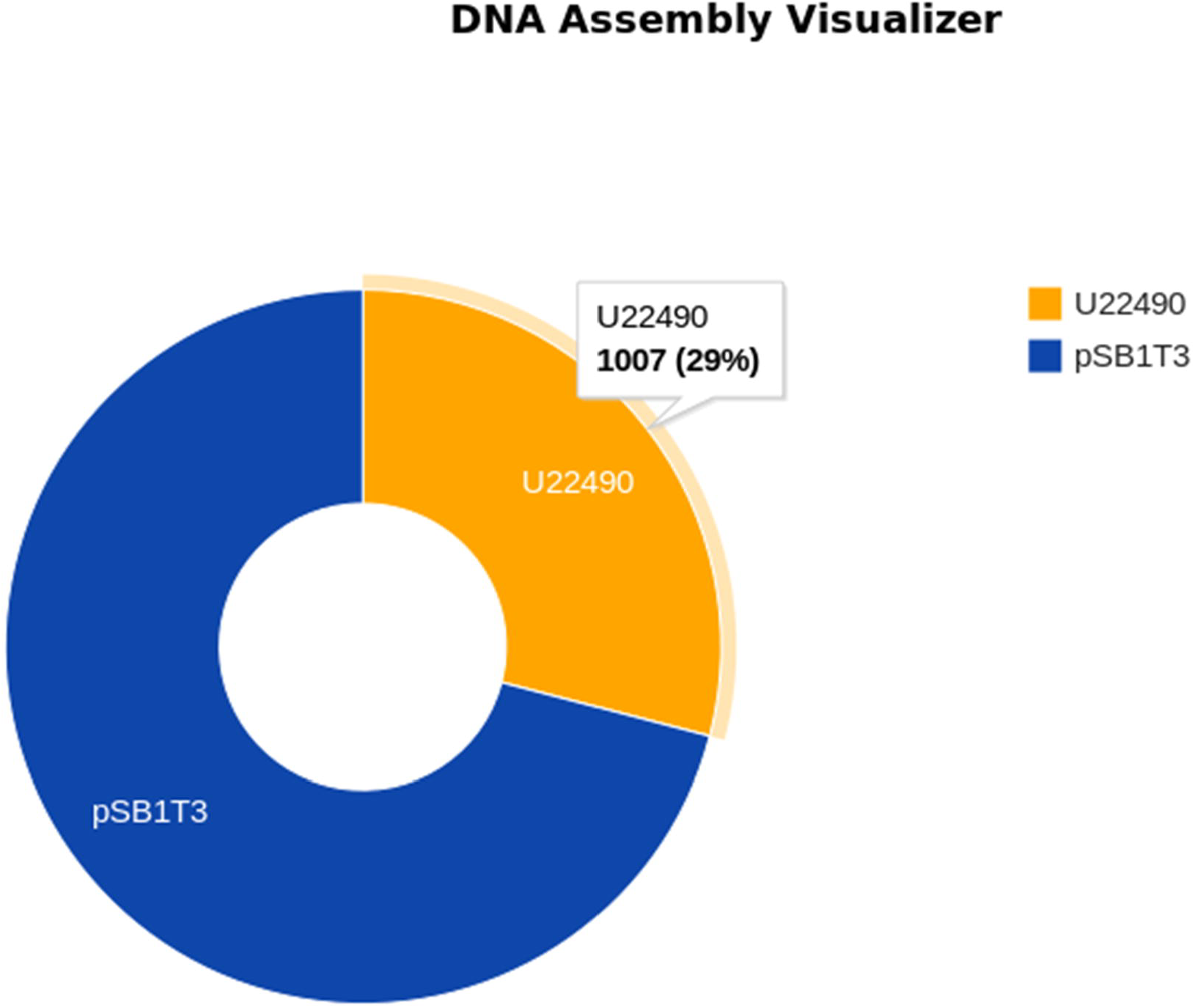
Assemblies’ visualization

### Results Validation

Once the previous stage was completed, test cases were created so that it included groups of possible paths between the Pyruvate and the Lactose, indicating the source maps used for the analysis. Afterward, these results were sent to the users that collaborated on the initial proposal of this research. Moreover, these users were also part of a biofuel production project, in which whey was transformed into biofuels through synthetic biology. The revision of the results was made by verifying the connectivity between the nodes, the subproducts produced by each reaction and the proteins involved in the process. From this, the corresponding observations were made, for the quality improvement and precision of the prototype’s results.

Some of the examples tested included the pathway generated from lactose to pyruvate via conventional glycolysis of D-glucose and the conversion of D-galactose to glycerol. Both pathways begin when a beta-galactosidase catalyzes the lactose in D-glucose and D-galactose. Then, D-glucose would follow the conventional glycolytic pathway while D-galactose would be transformed into glycerol via an alpha-galactosidase. However, the latter pathway did not take into account that the formation of glycerol is derived from the degradation of 3-beta-D-Galactosyl-sn-glycerol into glycerol and D-galactose and not a degradation product of D-galactose.

This error could be explained due to the reversibility of the reaction mediated by this enzyme since KEGG does not separate the direction of the reactions, causing the program to find a false link between these compounds. Other alternatives would be to select as final compound UDP-Glucose. In this case, the program should indicate the Leloir Pathway which begins with the enzyme GALM transforming β-D-galactose to α-D-galactose, followed by phosphorylation via a galactokinase and addition of an uridyl group via a galactose-1-phosphate uridyltransferase in order to obtain UDP galactose which would turn to UDP glucose via an epimerase. The program could also select dihydroxyace-tone phosphate as a final node, which should lead to a very different pathway involving phosphorylation of lactose (via PTS transport), breakdown into D-glucose and D-galactose-phosphate mediated by a phosphogalactosidase followed by conversion to tagatofuranose-6-phosphate by an isomerase which would be therefore phosphorylated forming D-tagatofuranose-1,6-bisphosphate. Finally, an aldolase would form dihydroxyacetone phosphate and D-glyceraldehyde 3-phosphate both of which can enter directly to glycolysis. In any case, the focus was to allow direct use of lactose into the carbon central metabolism leading, indirectly, to the increase of the production of fatty acids/triglycerides. Nevertheless, in order to achieve a major production of these compounds several physiological conditions should be required such as carbon: nitrogen ratio in media culture. To obtain better results and avoid this kind of errors, it would be necessary to change the interpretation of the KEGG pathways or use another database for which reversibility, or directed arcs, is expressed separately. Also, consideration of carbon loss or gain as well as reaction kinetics would considerably improve the applicability of this tool for its use in synthetic biology.

## 4 FUTURE WORK

It is important to mention that the restriction of entries allowed to establish a limited research scope; however, this does not restricts possible extensions of the tool:

1. *Change the input and output nodes:* although this research is mostly limited to Lactose and Pyruvate it is possible to use different compounds and maps, according to the end product that the user wants to get. However, it must be assured to select the proper metabolic pathways sample in order to have possible paths between the selected elements.
2. *Add new data repositories:* When including a new data source for metabolic pathways, it must be taken into account the connections to the protein and genes data bases. This allows for the maintenance of the relationship between metabolic pathways, protein and genes for the biobricks catalogue.
3. *Use several data repositories:* since each repository has its own output format, it is necessary to create a metabolic pathways translator to one only standard format that enables its manipulation as network structures.
4. *Validation of standards:* in terms of the tool release, it is important to review the standards in Bioinformatics & Nanobiotechnology for software development and genome editing (CRISPR).

## 5 CONCLUSIONS

Through the analysis of metabolic pathways of *E. coli* bacteria it was possible to design a tool for the purpose of assisting the biofuel production process. The translation of these pathways to graphs, using KEGG translator tool and well-known pathfinding algorithms such as Dijsktra and Breadth First Search, enable the creation and optimization of metabolic pathways. The analysis that was carried out over the existing maps were done to a depth of twelve elements and further.

Moreover, the implementation of a functional prototype for the design of this tool allowed for a more tangible validation in which pathways found will support the analysis and generation of new paths for the synthetic generation of biofuel. This was achieved through the transformation and analysis of more than one hundred metabolic pathways for *E. coli* bacteria, which were all the pathways that matched with the defined criteria, and the results integration with data repositories as KEGG, Uniprot and GenBank.

With these, a biobricks catalogue was created with the purpose of building new assemblies for the synthetic assembly of new organisms for biofuel production. The final product was characterized for its structure and functions. The structural characterization of the synthetic assemblies was obtained through the quantification of its nucleotide and amino acids numbers, which will enrich the design stage providing more information about the final result. The functional characterization of the assemblies was made based on the results obtained and the available data in UniProt, providing additional information about the involved proteins such as a basic description of its functions and the reactions that they catalyze.

Graphic representation techniques used on the prototype enabled the visualization of metabolic pathways through the JUNG framework and the final assemblies on pie charts, through Javascript web programming. This allows results to be shown in a simple way, making them more understandable for the user. The metabolic pathways transformations into graphs made the data analysis possible through the application of existing computing theories, such as data structure and search algorithms.

This design and prototype demonstrates the applicability of computing to a non-traditional command such as biology and computational biology, as well as the contribution to other fields such as synthetic biology and high performance computing, gene editing and supercomputing. The proposed solution enables the abstraction of organisms’ redesign process. It suggests possible optimized pathways between the present components in organic waste and biofuels, letting the user focus on problems of higher impact and complexity.

## ACKNOWLEDGEMENTS

To David García, Abad Rodríguez and Silver Ceballos for sharing their projects and knowledge in the field of synthetic biology, to Manuel Vargas for their guidance and suggestions for the project, to the National Council for Scientific and Technological Research in Costa Rica (CONICIT) and Intel Costa Rica for their financial support, to the Costa Rica Institute of Technology (TEC), University of Costa Rica (UCR) and LANOTEC for their support and collaboration to this project, to the provided computing services by cluster NELLY of Bioinformatics Medical School (UCR), to Antonio Solano for graphics elaboration and to everyone else that collaborated and supported this project: Carlos González, Diego Rivera-Gutiérrez, Carlos Álvarez, Mayra Rodríguez, Shannon Cummings, María Catalina Rosales, Juan Carlos Saborío-Morales, María Mora, Mayela Guzmán, Marianela Snowball, Marco Loaiza, Karen Rodríguez and Adalberto Rodríguez.

## Notes

https://gitlab.com/lvasquezcr/pathways-analyzer

